# Mutation of the *P. falciparum* flavokinase confers resistance to roseoflavin and 8-aminoriboflavin

**DOI:** 10.1101/2024.04.04.588205

**Authors:** Ayman Hemasa, Christina Spry, Matthias Mack, Kevin J. Saliba

## Abstract

We previously found that two riboflavin analogues, roseoflavin and 8-aminoriboflavin, inhibit malaria parasite proliferation by targeting riboflavin utilisation. To determine the mechanism of action of roseoflavin in *P. falciparum*, we generated roseoflavin-resistant parasites by *in vitro* evolution over 27 weeks. The roseoflavin-resistant parasites were found to be four times more resistant to roseoflavin and cross-resistant to 8-aminoriboflavin. Resistant parasites were subjected to whole genome sequencing and a missense mutation (T2015A), leading to an amino acid exchange (L672H), was detected in the gene coding for a putative flavokinase (*Pf*FK), the enzyme responsible for converting riboflavin (vitamin B_2_) into the cofactor flavin mononucleotide (FMN). To confirm that the L672H mutation is responsible for the observed phenotype, we generated parasites with the missense mutation incorporated into the *Pf*FK gene *via* a single-crossover recombination. The IC_50_ values for roseoflavin (RoF) and 8-aminoriboflavin against the RoF-resistant parasites created through *in vitro* evolution were indistinguishable from the IC_50_ values for parasites in which the missense mutation was specifically introduced into the native *Pf*FK. To investigate this mutation, we generated two parasite lines episomally-expressing GFP-tagged versions of either the wild type or mutant forms of flavokinase. We found that *Pf*FK-GFP localises to the parasite cytosol and that immunopurified *Pf*FK-GFP was active and phosphorylated riboflavin into flavin mononucleotide. The L672H mutation caused a reduction of the binding affinity, especially for the substrate RoF, which explains the resistance phenotype. The mutant *Pf*FK is no longer capable of phosphorylating 8-aminoriboflavin, but its antiplasmodial activity against resistant parasites can still be antagonised by increasing the extracellular concentration of riboflavin, consistent with the compound also inhibiting parasite growth through competitive inhibition of *Pf*FK. Our findings, therefore are consistent with roseoflavin and 8-aminoriboflavin inhibiting parasite growth by inhibiting FMN production, in addition to the generation of toxic flavin cofactor analogues.

## Introduction

Malaria is a disease caused by unicellular protozoan parasites of the genus *Plasmodium.* Six species infect humans ^1^. The parasite is spread by female *Anopheles* mosquitoes, infecting an estimated 249 million estimated cases in 2022 and resulting in the death of 608,000 people ^2^. To combat the rising incidence of *Plasmodium falciparum* (the most virulent *Plasmodium* species infecting humans) resistant to antimalarials, including artemisinin-based combination therapy, regarded as the frontline treatment for uncomplicated malaria ^3^, new antimalarial drugs and drug targets are required ^4^.

Understanding the nutritional requirements of the intraerythrocytic stage of the malaria parasite could reveal critical mechanisms utilised by the parasites that may be exploited for the development of new antimalarials. Although we have gained some knowledge of the parasite’s need for specific vitamins, such as pantothenate ^5, 6^ and thiamine ^7–9^, we still understand very little about the parasite’s need for other vitamins. Riboflavin (vitamin B_2_) is the essential precursor of two important enzyme cofactors – flavin mononucleotide (FMN) and flavin-adenine dinucleotide (FAD; **Figure 1**). FMN and FAD are used as cofactors by many enzymes ^10^. For example, the *P. falciparum* dihydroorotate dehydrogenase (*Pf*DHODH) is an FMN-dependent enzyme that is critical for the parasite’s pyrimidine production ^11^. Yeast, plants, and most prokaryotes can synthesize riboflavin *de novo* ^12^, whereas many animals, including humans, lack this ability and rely on the uptake of riboflavin from external sources ^13^. *P. falciparum* parasites are auxotrophic for riboflavin, believed to obtain riboflavin from their host cells ^14, 15^.

**Figure 1:**
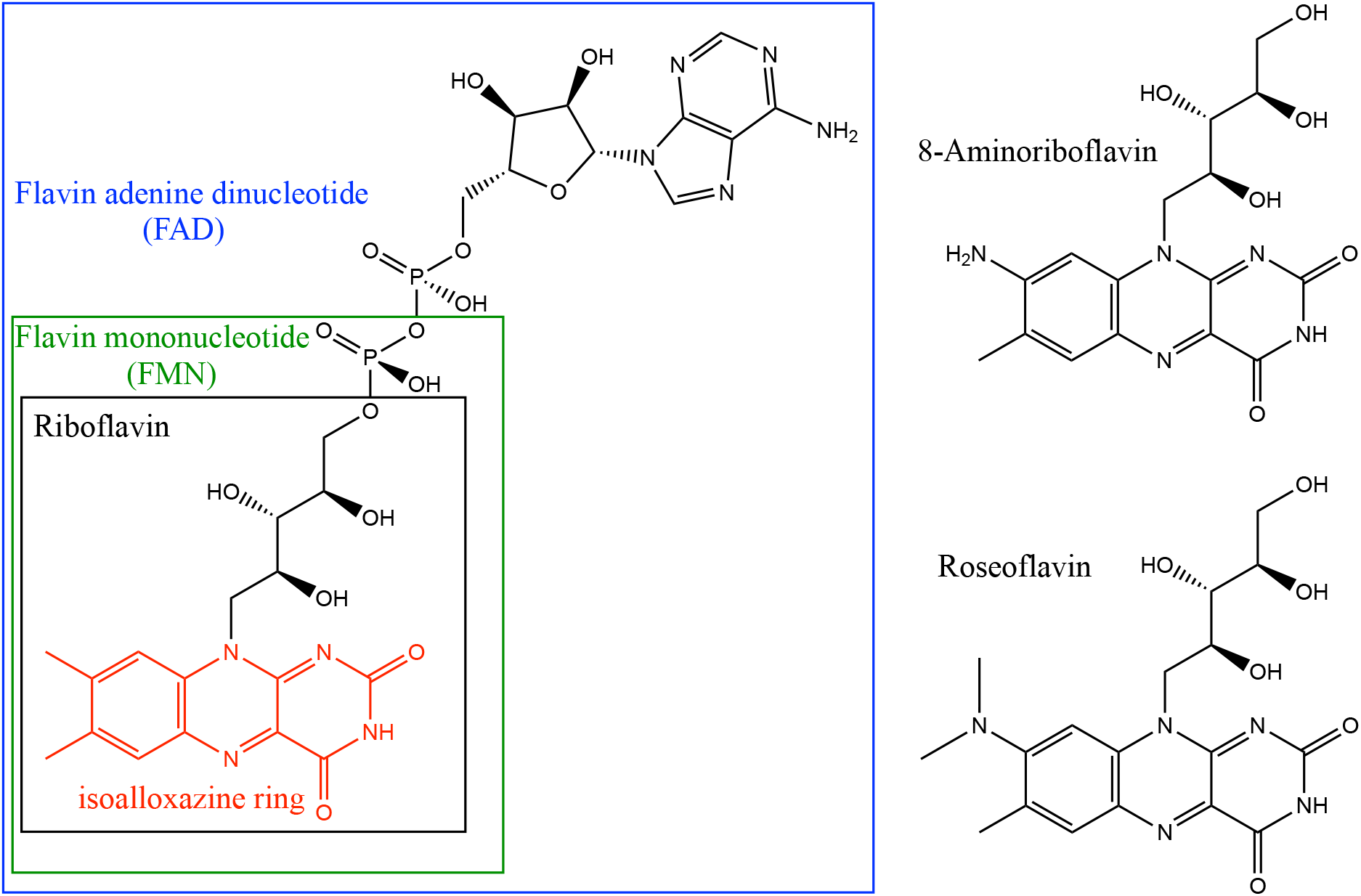
Structure of riboflavin (black box) FMN (green box) and FAD (blue box) with the oxidized form of the 7,8-dimethyl-isoalloxazine moiety (red structure). The structures of roseoflavin and 8-aminoriboflavin are included (outside of the blue box) for comparison.

The synthesis of FMN and FAD requires two enzymatic steps. First, riboflavin is phosphorylated by flavokinase (riboflavin kinase) to form FMN, a redox-active cofactor. Subsequently, FAD-synthetase adenylylates FMN to produce FAD. Both steps are ATP dependent.

We have recently shown that two analogues of riboflavin, namely roseoflavin (RoF, **Figure 1**) and 8-aminoriboflavin (8AF, **Figure 1**), inhibit the *in vitro* proliferation of *P. falciparum* by interfering with the utilisation of riboflavin ^16^. We also found that RoF and 8AF reduce FMN and/or FAD levels within *P. falciparum* parasites, consistent with RoF and 8AF interacting with riboflavin metabolism enzymes, either as substrates or inhibitors ^16^. However, the exact mechanism of action of these compounds is unknown.

In this study, we have used *in vitro* evolution to generate RoF resistant *P. falciparum* parasites with the objective of understanding the mode of action of RoF and 8AF. Our results are consistent with the antiplasmodial activity of the riboflavin analogues being due to both a reduction of FMN levels (by competing with riboflavin for *Pf*FK binding) as well as generation of toxic flavin cofactor analogues.

## Methods

### Cell Culture and Media

*P. falciparum* (3D7 strain) parasites were maintained in culture within human erythrocytes as previously described ^17, 18^. RPMI-1640 culture medium was supplemented with 200 µM hypoxanthine, 24 mg/L gentamycin, 11 mM D-glucose and Albumax II (0.6%, w/v). Riboflavin-free RPMI-1640 (R8999-14, USBiological Life Sciences, USA, which is also free of folic acid and L-glutamine) was prepared by dissolving 10.092 g in 1 L water, supplementing with 2.26 µM folic acid, 2.054 mM L-glutamine, adjusting to pH 7.4 and filter sterilising. The riboflavin-free medium was then further supplemented with the same additives as RPMI-1640 culture medium. Parasites were maintained at 4% hematocrit (HCT) in O^+^ erythrocytes, flushed with a gas mixture of 3% CO_2_, 1% O_2_ and 96% N_2_, and maintained at 37°C in a horizontal shaker. Every 24 h, the suspension was centrifuged at 500 × *g* for 5 min. The supernatant was replaced with fresh medium, and the infected erythrocytes diluted 10-20 times with uninfected erythrocytes when the parasites were in the trophozoite stage. The parasitaemia was maintained within the range of 1-6%.

### *In vitro* antiplasmodial growth assays

The antiplasmodial activity assays were performed in 96-well plates using the Malstat or SYBR-safe assay, as previously described ^19–21^, with minor modifications ^16^. At the end of the compound incubation period, plates were stored at −20 °C for at least 24 hours prior to processing using SYBR-safe lysis or Malstat solutions. A Fluostar OPTIMA multi-detection microplate reader with excitation and emission wavelengths of 490 and 520 nm, respectively, was used to detect the fluorescence in the SYBR-safe assay. For the Malstat assay, the absorbance was measured at 620 nm using the same plate reader. The 50% inhibitory (IC_50_) values of test compounds were determined by fitting the data to a sigmoidal curve using nonlinear least squares regression.

### *In vitro* evolution and whole genome sequencing

Two independent 10 mL cultures of synchronised ring-stage *P. falciparum* parasites (from a frozen stock of a previously re-cloned 3D7 line ^22^) at 2-3% parasitemia and 2% HCT, were exposed to RoF at 1.6 µM (the IC50 value against the parental line in the presence of the standard concentration of riboflavin present in RPMI-1640). In the experiment, RoF concentrations were increased gradually over 27 weeks until the IC50 of RoF became four times higher than the initial control value. After maintaining the parasites in culture for four weeks without RoF, we confirmed that the sensitivity of RoF-resistant parasites had not changed during the RoF-free culture period. Four random clones (see below) were then selected for further study. These clones along with the parent line were subjected to whole genome sequencing (WGS) using the Illumina MiSeq platform (paired-end reads, 2 × 250 base pairs) Nextera XT Kit. To detect any mutations, PlaTyPus, an integrated variant pipeline, was used to analyse the WGS data, with minor modifications ^23^.

### Cloning of *P. falciparum* parasites and parasite enrichment

Cloning was performed by either limiting dilution in a 96-well plate at a density of 0.5 parasites *per* well ^24^ or by delivering one infected erythrocyte per well, into a 96-well plate using flow cytometry ^25^. For cloning using flow cytometry, infected erythrocytes were enriched to a parasitaemia >95% as described previously ^26^. Erythrocytes infected with parasites expressing a GFP-tagged protein were not enriched, but were instead sorted on the basis of GFP fluorescence. After 3 weeks of incubation, wells were inspected for the presence of parasites using either Giemsa-stained blood smears or Malstat reagent ^20, 21^.

### Construction of plasmids and purification of flavokinase

For expression of WT and mutant forms of *P. falciparum* flavokinase (*Pf*FK) as a GFP fusion protein, the coding sequence of *Pf*FK was amplified from the gDNA of wild-type and RoF-resistant *P. falciparum* parasites, respectively, employing the following forward and the reverse primers, 5′-GGAATTG<u>CTCGAG</u>ATGGTTCATGATAAATATCATAAAATAGC-3′ and 5’-CACT<u>GGTACC</u>TTTTATATTTTCGAGAAAATACTTAC-3′. The restriction endonuclease sites, XhoI and KpnI, are underlined. The XhoI/KpnI-treated PCR product, of the WT and mutant *Pf*FK gene, were ligated to the XhoI/KpnI digested pGlux-1 plasmid (provided by Prof Alexander Maier, Australian National University) which confers ampicillin resistance. The WT and mutant-*Pf*FK-pGlux-1 constructs were transformed into RbCl competent *E. coli* (DH5α). The construct was then purified using a Qiagen HiSpeed Plasmid Maxi Kit and 100 µg of the purified construct transfected into ring-stage *P. falciparum* parasites. Successfully-transfected parasites were selected with WR99210 (10 nM) for three weeks ^27^. Both WT and mutant *Pf*FK-GFP were purified by immunoprecipitation using GFP-Trap ^28, 29^ from saponin-isolated ^5^, trophozoite-stage parasites. The purified proteins from 1 × 10^10^ isolated trophozoites were immediately used in enzyme assays and western blots.

### Enzyme assays and HPLC analysis of flavins

Enzyme activity of the purified WT and mutant *Pf*FK was measured in 1 mL of 50 mM potassium phosphate (pH 7.5), containing 0.5-200 µM of the test substrates, riboflavin, RoF or 8AF, 3 mM ATP, 24 mM sodium dithionite (Na_2_S_2_O_4_), 12 mM MgCl_2_, and 50 µL of the purified *Pf*FK-GFP, as previously described for flavokinase enzymes from other organisms, with some modifications ^30^. The reaction was carried out at 37 °C. The samples were agitated at 1500 rpm for the duration of the experiments using an Eppendorf Thermomixer to keep the enzyme (which is attached to the anti-GFP-beads) suspended during the reaction. Reactions were initiated by the addition of *Pf*FK-GFP and monitored for up to 3 h. At appropriate time intervals, 180 µL aliquots were removed and 1.8 µL of trichloroacetic acid (TCA) added to a final concentration of 1% (v/v) to stop the reaction. The aliquots were then filtered using nylon syringe filters (0.45 µm pore size, 13 mm diameter) into micro-sampling, amber glass HPLC vials (vial volume capacity was 300 µL), and 45 µL applied directly to an HPLC column (Kinetex® 2.6 µm (particle size) Polar C18 100 Å (pore size), 30 mm (length) × 2.1 mm (internal diameter)). The following solvent system was used at a flow rate of 1.2 mL/min: 20 mM potassium phosphate pH 3.5 (mobile phase A), 100% methanol (mobile phase B). The entire duration of the run is 16 minutes, during which the composition consists of 100% phase A and 0% phase B for the initial 12 minutes. Subsequently, a gradient of 40% phase A and 60% methanol is employed from 10 to 12 minutes to eliminate nonpolar impurities. The system is then reverted to 100% phase A from 12 to 16 minutes, concluding the run. Detection of RoF and RoFMN (phosphorylated RoF, an FMN analogue) was carried out using a diode array detector (absorbance at 503 nm), whereas the detection of riboflavin, FMN, 8AF and 8AFMN (phosphorylated 8AF, an FMN analogue) was performed using a Dionex fluorescence detector (excitation at 480 nm and emission at 520 nm). The peak area of the metabolites was quantified using standard curves. Flavokinase activity was expressed as a function of the amount of metabolite generated (in micromolar) from the corresponding substrate *per* 10^13^ cells *per* hour. All K_m_ and V_max_ values were determined from fitted Michaelis-Menten curves.

### Western blots

Denaturing western blots were performed with saponin-isolated parasite lysates and immunoprecipitated *Pf*FK-GFP. Protein samples were separated using polyacrylamide gel electrophoresis (PAGE) in NuPAGE (4–12%) gels (Life Technologies), then transferred to nitrocellulose membranes and blocked in 4% w/v skim milk powder in Phosphate Buffered Saline (PBS). Blocked membranes were subjected, for 2 h, to anti-GFP mouse antibody 400 µg/mL (primary antibody) and goat anti-rabbit HRP (secondary antibody). Membranes were then incubated in Pierce Enhanced Chemiluminescence (ECL) Plus Substrate (Life Technologies) according to the manufacturer’s specifications and protein bands visualised on a ChemiDoc MP Imaging System.

### Introduction of L672H mutation into the endogenous *Pf*FK

To generate a mutant *Pf*FK-GFP_glmS construct, a C-terminal homologous region (HR, 774 base pairs of the *Pf*FK gene including the sequence encoding the L672H mutation) was amplified using the mutant *Pf*FK gDNA and the forward primer, 5′-GC<u>ACTAGT</u>AT**A**TCTCA**T**GCTAA**G**AGACATGGTA-3′ (3 silent mutations included and in bolded letters), and the reverse primer 5′-GGAATTG<u>CTCGAG</u> TTATTTTATATTTTCGAGAAAATACTTACATTGTTGAAAAGTTTCATTATTTTTCAATTTGTTAAGTACAATTCTGGCTAGTTCACAATCAAATTGAATAGCTTGAATATGATG-3′, cloned into eGFP-glmS plasmid (provided by Parichat Prommana, Thailand, ^31^). The restriction endonuclease sites SpeI and XhoI are underlined. The resulting PCR product was ligated into the corresponding SpeI/XhoI-digested eGFP-glms plasmid. A specific guide RNA sequence was selected 774 bp upstream from the C-terminus of *Pf*FK and generated by annealing two overlapping oligonucleotides having overhangs complimentary to the BbsI sites. The sequence of these two oligonucleotides were: 5′-<u>TATT</u>ATTTCTCACGCTAAAAGACA-3′, 5′-<u>AAAC</u>TGTCTTTTAGCGTGAGAAAT-3′ (overhangs are underlined). The annealed gRNA product was inserted into BbsI-digested pDC2-cam-coCas9-U6-hDHFR vector containing a codon-optimised (co) Cas9 driven by the *Pf-*calmodulin promoter, and a U6 cassette for gRNA expression (construct provided by Dr Marcus Lee, Hinxton, UK ^32^. Parasite transfection was carried out simultaneously with these two constructs, *Pf*FK (homologous region of 774 base pair, m-HR)_GFP_glmS and gRNA-*Pf*FK-pDC2-cam-coCas9-U6-hDHFR. Transfectants were obtained following four weeks of selection with 10 nM WR99210 and 2 µg/mL blasticidin. Integration of the mutant HR of *Pf*FK-GFP_glmS was confirmed after three blasticidin on/off cycles of three weeks duration each, followed by magnetic parasite enrichment and then parasite cloning using FACS into 96 well plate (one parasite per well) ^33^. Successful integration of the construct was confirmed by PCR.

### Fluorescence microscopy

Fluorescence microscopy was carried out on a Deltavision Deconvolution microscope (100× oil objective with a resolution of 0.066 µm per pixel). Parasite DNA was stained with Hoechst 33342 (4 µg/mL for 15 min). Images were collected at ambient temperature, deconvoluted and linearly adjusted for contrast and brightness.

### Flow cytometry

Flow cytometry was used to quantify the fraction of GFP-positive parasites (i.e. parasites expressing *Pf*FK-GFP). Mid-trophozoite-stage cultures expressing GFP-tagged flavokinase (either episomally or endogenously) were preincubated with 4 µg/mL of Hoechst stain, to distinguish the iRBCs from the uRBCs, for 15 min followed by washing the cells three times with PBS. Aliquots of 10-20 µL from all samples were diluted in 200-300 µL PBS to a concentration of 10^5^-10^6^ cells per mL and applied to 1.2 mL Costar polypropylene cluster tubes (Corning) and subjected to flow cytometry. Non-GFP-expressing 3D7 WT parasites were used to gate non-fluorescent parasites. Data were analysed using BD FACSDiva software © Becton, Dickinson and Company (Forward scatter 450 V, Side scatter 350 V and Alexa Fluor 488 = 600 V, Pacific Blue, 450/50 nm).

### Alignment of flavokinases

Using PROMALS3D ^34^, the flavokinase gene from *P. falciparum* was aligned with other flavokinases from various organisms. The following flavokinase homologues were aligned (accession number included in brackets): *P. falciparum* (Q8IDB3); *Homo sapiens* (Q969G6); *Schizosaccharomyces pombe* (O74866); *Bacillus subtilis* (P54575); *Trypanosoma brucei brucei* (Q38DG4); *Arabidopsis thaliana* (Q84MD8); *Streptococcus agalactiae serotype* Ⅲ (Q8E5J7); *Candida albicans* (Q5A015); *Trichophyton rubrum* (F2SJS4); *Saccharomyces cerevisiae* (Q03778); *Corynebacterium ammoniagenes* (Q59263).

### Statistical analysis

Statistical analysis was carried out with unpaired, two-tailed, Student’s t-test and ordinary one-way ANOVA using GraphPad Prism 9.3 (GraphPad Software, Inc) from which the 95% confidence interval of the difference between the means (95% CI) was obtained. GraphPad Prism 9.3 version for Windows 11 was used for all regression analysis.

## Results

### *In vitro* evolution of *P. falciparum* in the presence of RoF resulted in parasites resistant to RoF and 8AF that harboured a mutation in *Pf*FK

Using a cloned 3D7 *P. falciparum* parasite line ^22^, two independent cultures were exposed to continuous drug pressure with increasing concentrations of RoF, starting with the IC_50_ value (1.6 µM). When the sensitivity of the parasites to RoF decreased by approximately 4-fold (following 27 weeks of drug pressuring; IC_50_ = 5.6 ± 0.3 µM), they were maintained in the absence of RoF for 4 weeks. The parasites remained resistant to RoF during this period (**Figure 2A**). The parasites were then cloned by limiting dilution and four parasite clones established, two from each independent *in vitro* evolution experiment (RoF-A-E8, RoF-A-B6, RoF-B-E11, RoF-B-G2). The IC_50_ values (means ± SEM) of RoF against RoF-A-E8, RoF-A-B6, RoF-B-E11, RoF-B-G2 were 4.4 ± 0.3-fold (6.4 ± 1.5 µM, 6.1 ± 1.6 µM, 8.1 ± 2.3 µM, 7.5 ± 2.5 µM, respectively) higher than the IC_50_ against the parent parasites (IC_50_ = 1.6 ± 0.1 µM, P<0.05) (**Figure 2A**). To investigate whether the RoF resistant parasites were cross-resistant to 8AF, we determined the sensitivity of the resistant parasites to 8AF. As the solubility of 8AF only permits testing up to 50 µM, we assessed the sensitivity in riboflavin-free RPMI where the IC_50_ against WT parasites is 2 ± 0.4 nM rather than complete RPMI where the IC_50_ of 8AF is 7 ± 1 µM ^16^. The IC_50_ values of 8AF in riboflavin-free medium against the RoF-A-E8, RoF-A-B6, RoF-B-E11, RoF-B-G2 parasite lines (1.2 ± 0.1 µM, 1.1 ± 0.1 µM, 1.2 ± 0.1 µM, 1.1 ± 0.1 µM, respectively) were found to be 1000-fold higher than those of the parent strain (0.0010 ± 0.0002 µM, P<0.0001, unpaired t-test; **Figure 2B**). To eliminate the possibility that these clones have developed a non-specific resistance mechanism, we tested the sensitivity of the clones to chloroquine, an antiplasmodial with a mechanism of action that is unrelated to riboflavin metabolism and/or utilisation ^35^. We found that the chloroquine IC_50_ values against the RoF-resistant parasites were not statistically different from those of the parent strain (P = 0.44, unpaired t-test) (**Figure 2C**).

**Figure 2:**
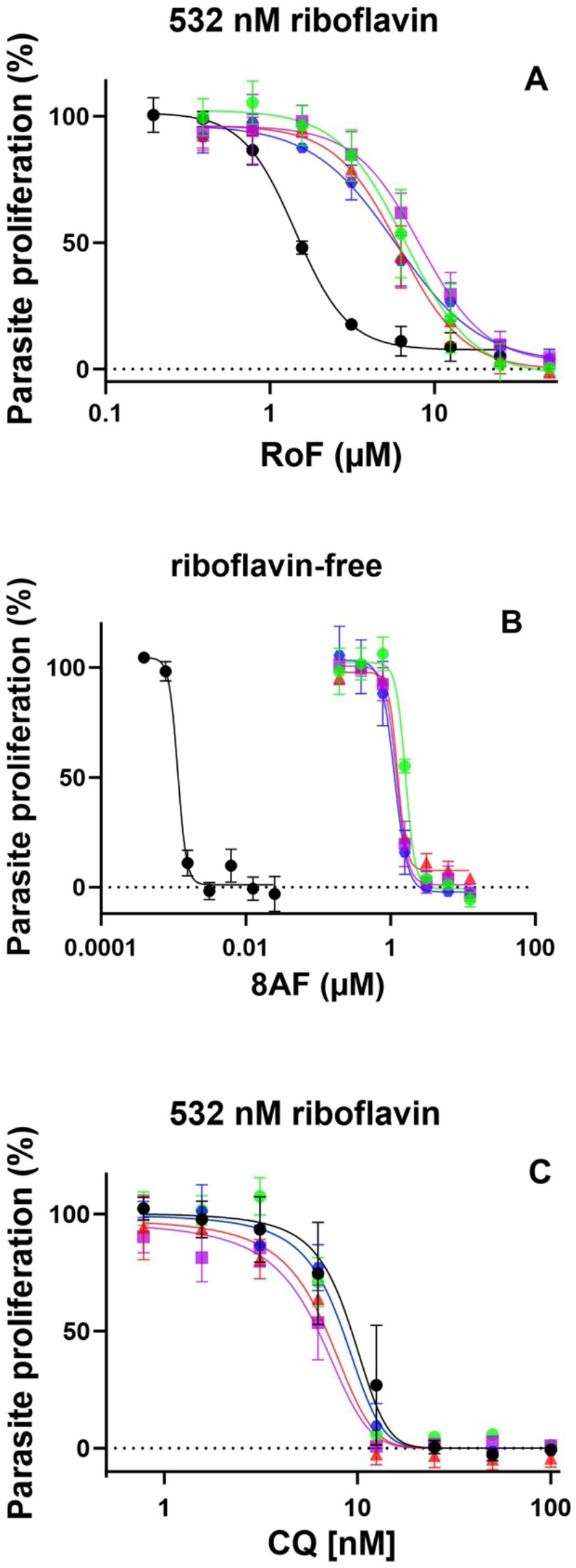
Percentage parasite proliferation of the parent line (black circles) and the four RoF-resistant clones, RoF-A-E8 (red triangles), RoF-A-B6 (blue hexagon), RoF-B-E11 (purple squares), RoF-B-G2 (green circles) in the presence of (A) RoF in normal RPMI-1640 (containing 532 nM riboflavin), (B) 8AF in riboflavin-free RPMI-1640 or (C) chloroquine in normal RPMI-1640. Data points are averaged from 3-4 independent experiments, each carried out in triplicate. Error bars represent SEM and are not visible if smaller than the symbols.

To determine the mutation/s responsible for the RoF and 8AF resistance phenotype, gDNA from each clone was extracted and subjected to whole genome sequencing. We found that all four clones harbour two non-synonymous mutations, one in a gene coding for a putative riboflavin kinase, *Pf*FK (PF3D7_1359100), L672H, and the other in the gene coding for an apicoplast ribosomal protein S5 (*PF*C10_API0033), R118K, **Table S1**). Several other non-synonymous mutations were also detected across the clones, but none of these mutations occurred in all of the clones (**Table S1**). A structure-based alignment of the *Pf*FK sequence with that of flavokinases from bacteria (*B. subtilis*, *S. agalactiae*, *C. ammoniagenes*), yeast (*S. pombe*, *C. albicans* and *S. cerevisiae*), the fungus *T. rubrum*, the plant *A. thaliana*, the parasite *T. brucei*, and *H. sapiens*, for which crystal structures are available, showed that L672 is a highly conserved residue (100% identity; **Figure S1**). In the absence of 3D structures for *Pf*FK, we aligned an AlphaFold-predicted *Pf*FK structure (AF-Q8IDB3-F1, ^36^ with the human flavokinase, for which a crystal structure is available (**Figure 3**). L672 is in a region of *Pf*FK that is predicted with high confidence (per-residue Local Distance Difference Test, pLDDT >70 but <90). The residue is within an alpha-helix in a position overlapping with the corresponding residue in human flavokinase (L115, **Figure 3**). In the human flavokinase, L115 forms part of the riboflavin or FMN binding site, with the side chain of L115 forming hydrophobic interactions with the isoalloxazine ring (**Figure 3**). Mutation of L672, a nonpolar amino acid, to a polar histidine residue, which can be positively charged at physiological pH, is likely to disrupt the hydrophobic interactions with riboflavin.

**Figure 3:**
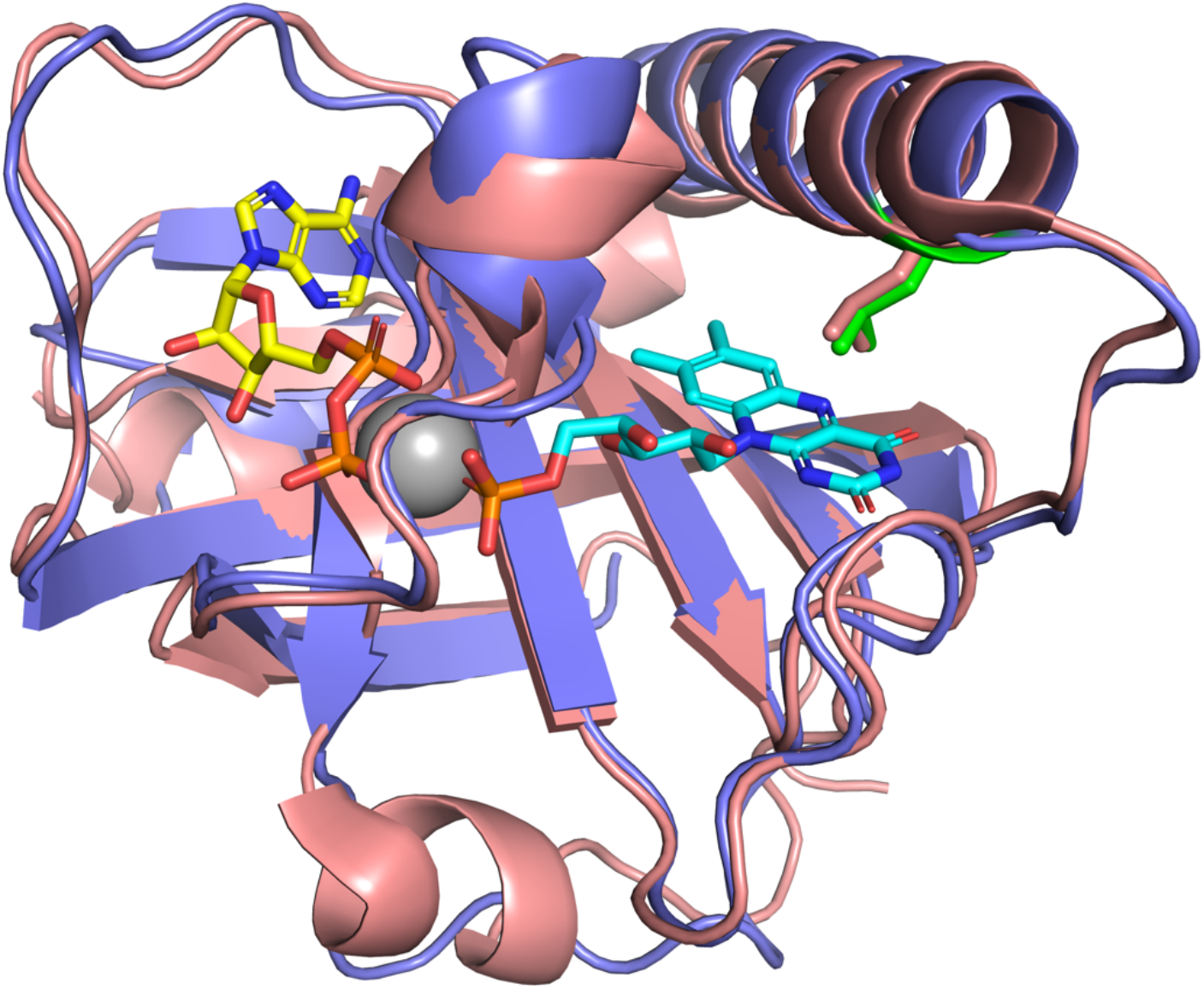
Overlay of the predicted three-dimensional structure of the C-terminal domain of *Pf*FK (Y565-K707, purple, predicted using AlphaFold, AF-Q8IDB3-F1) with the experimentally determined X-ray crystal structure of the entire (151 amino acid) human flavokinase (pink) with FMN (carbons in blue), ADP (carbons in yellow) and Mg^2+^ (grey) bound (PDB ID: 1Q9S). The side chains of L672 (green) and the corresponding residue in the human flavokinase (L115, pink) are shown. *Pf*FK shares 40% sequence identity with human flavokinase in this region. For clarity, residues M1-K564 of *Pf*FK is not included in the figure.

### L672H mutation is responsible for the resistance phenotype against RoF and 8AF

To determine whether the L672H mutation is responsible for the resistance phenotype, we generated transgenic 3D7 *P. falciparum* parasites that, in addition to expressing the endogenous *Pf*FK, also episomally express a GFP-tagged version of the wild-type *Pf*FK or a copy of *Pf*FK harbouring the L672H mutation. We found that the sensitivity to RoF and 8AF of parasites expressing only the wild-type form of *Pf*FK (i.e. the endogenous version in addition to the GFP-tagged episomally-expressed version) did not change when compared to the sensitivity of the parent parasite line (P = 0.61 and 0.22 unpaired t-test, N = 3, respectively, **Figure 4A** and **4B**). In contrast, we found that parasites episomally expressing the mutant form of *Pf*FK tagged to GFP (in addition to expressing the endogenous, wild-type version) were 3-times more resistant to RoF (IC_50_ = 4.2 ± 0.4 µM; **Figure 4A**), and 2.7-times more resistant to 8AF (IC_50_ = 19 ± 3 µM; **Figure 4B**), in normal RPMI-1640 medium, when compared to wild-type parasites (IC_50_ = 1.4 ± 0.3 and 7 ± 1 µM, P = 0.005 and 0.021, respectively) and parasites additionally expressing the wild-type form of *Pf*FK tagged to GFP (IC_50_ = 1.6 ± 0.1 and 9 ± 1 µM, P = 0.007 and 0.017, respectively; **Figure 4**). The IC_50_ of RoF against roseoflavin-resistant parasites (8.1 ± 2.3 µM) was found to be 2-fold higher than the IC_50_ of RoF against the parasites episomally expressing the mutant form of *Pf*FK (4.2 ± 0.4 µM), **Figure 4A**). It is clear that the RoF-resistant parasites are also less sensitive to 8AF than parasites episomally expressing the mutant form of *Pf*FK **(Figure 4B)**. However, the precise fold difference in sensitivity in normal RPMI medium (where the riboflavin concentration is 0.532µM) could not be determined because at 25 µM (the highest concentration that can be tested due to limited solubility) there was less than 50% inhibition. However, by extrapolating the curve, an IC_50_ value of 70 µM was estimated for 8AF against these parasites; 3.7-fold higher than the 8AF IC_50_ observed against parasites episomally expressing the mutant form of *Pf*FK (19 ± 3 µM).

**Figure 4:**
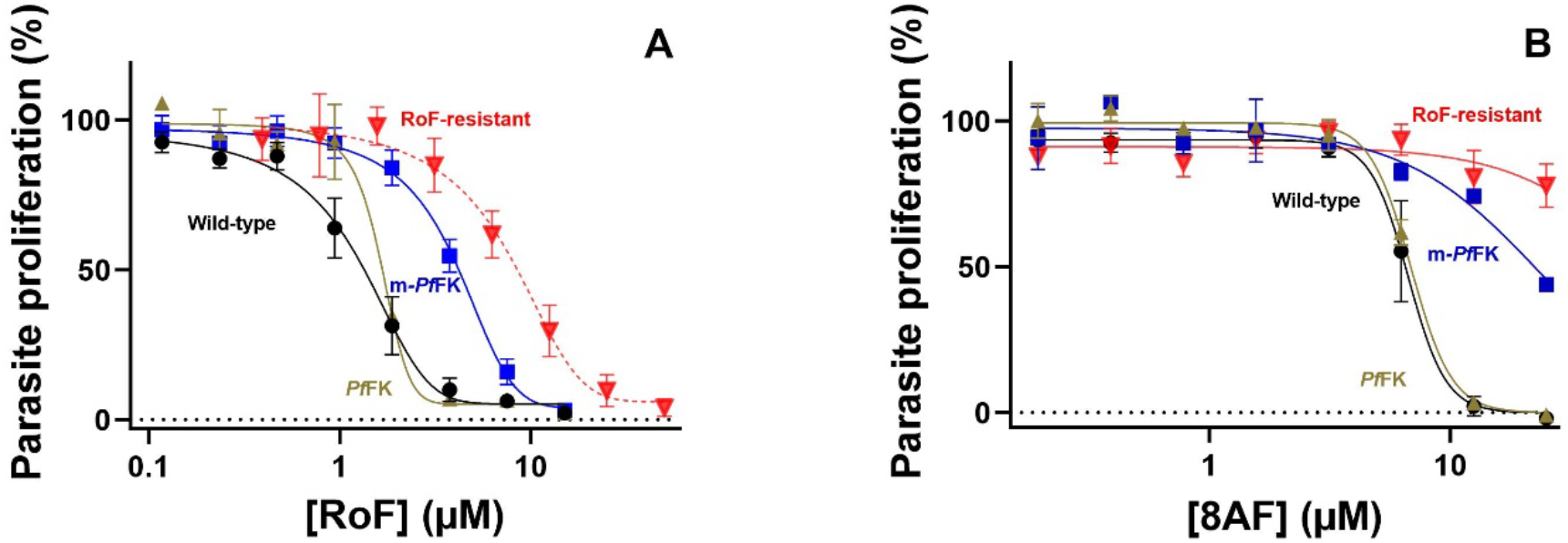
Proliferation of wild-type parasites (black circles), parasites episomally expressing the wild-type form of *Pf*FK tagged to GFP (gold triangles), parasites episomally expressing mutant *Pf*FK tagged to GFP (blue squares), and RoF-B-E11 (red inverted triangles) in the presence of (A) RoF and (B) 8AF in normal RPMI-1640. The RoF-B-E11 data in A (dotted line) is the same data set as that shown in Figure 2B and has been included here for comparison. Parasite lines episomally expressing a copy of *Pf*FK-GFP (wild-type or mutant) also express the endogenous wild-type copy of *Pf*FK. Values are averaged from three independent experiments, each carried out in triplicate. Error bars represent SEM and are not visible if smaller than the symbols.

These data are consistent with the L672H mutation being associated with the resistance phenotype. It is also evident that the episomal expression of the mutant *Pf*FK in transgenic parasites, alongside the endogenous, wild-type *Pf*FK, is not sufficient to confer resistance to the level observed in RoF-B-E11. To explore more definitively whether the L672H mutation is responsible for the resistance mechanism, we introduced the L672H mutation in the endogenous *Pf*FK gene.

### Single-crossover-mediated integration of the L672H mutation confers resistance to RoF and 8AF

We attempted CRISPR/cas9 editing of the *Pf*FK gene to introduce the T2015A mutation (resulting in the L672H mutation in the protein) into the endogenous gene. Initially, only a small proportion of parasites appeared to have integrated the mutated *Pf*FK gene construct, as judged by PCR. Uniform integration was obtained following three on/off cycles, with each off cycle lasting three weeks, of selection with 2 µg/mL blasticidin. This was then followed by FACS-mediated cloning. Sequencing of the *Pf*FK gene region of three of the clones showed that the integration had occurred *via* a single crossover recombination event rather than CRISPR/cas9. Parasites harbouring the single-crossover-mediated L672H mutation in *Pf*FK were then tested for their sensitivity towards RoF and 8AF. We found these parasites to be resistant to RoF by 4.5-fold in normal RPMI-1640 (IC_50_ = 4.1 ± 0.1 µM, P <0.0001, unpaired, two tailed t-test, N = 5, **Figure 5A**) and 500-fold resistant to 8AF in riboflavin-free RPMI-1640 (IC_50_ = 0.5 ± 0.1 µM, P = 0.0019, unpaired, two tailed t-test, N =3, **Figure 5B)**, as determined by comparison with the parent parasites in the corresponding RPMI-1640 (IC_50_ = 0.9 ± 0.2 µM for RoF and 0.001 ± 0.002 µM for 8AF, **Figure 5A** and **5B**). There was no significant difference in the IC_50_ values of RoF and 8AF against RoF-resistant parasites generated *via in vitro* evolution (4.0 ± 0.3 and 0.5 ± 0.1 µM, respectively), and parasites that had the L672H mutation selectively integrated into the endogenous *Pf*FK (4.1 ± 0.1 and 0.5 ± 0.1 µM, P = 0.73 and 0.99, respectively, unpaired, two tailed t-tests, N = 3, **Figure 5A** and **5B**).

**Figure 5:**
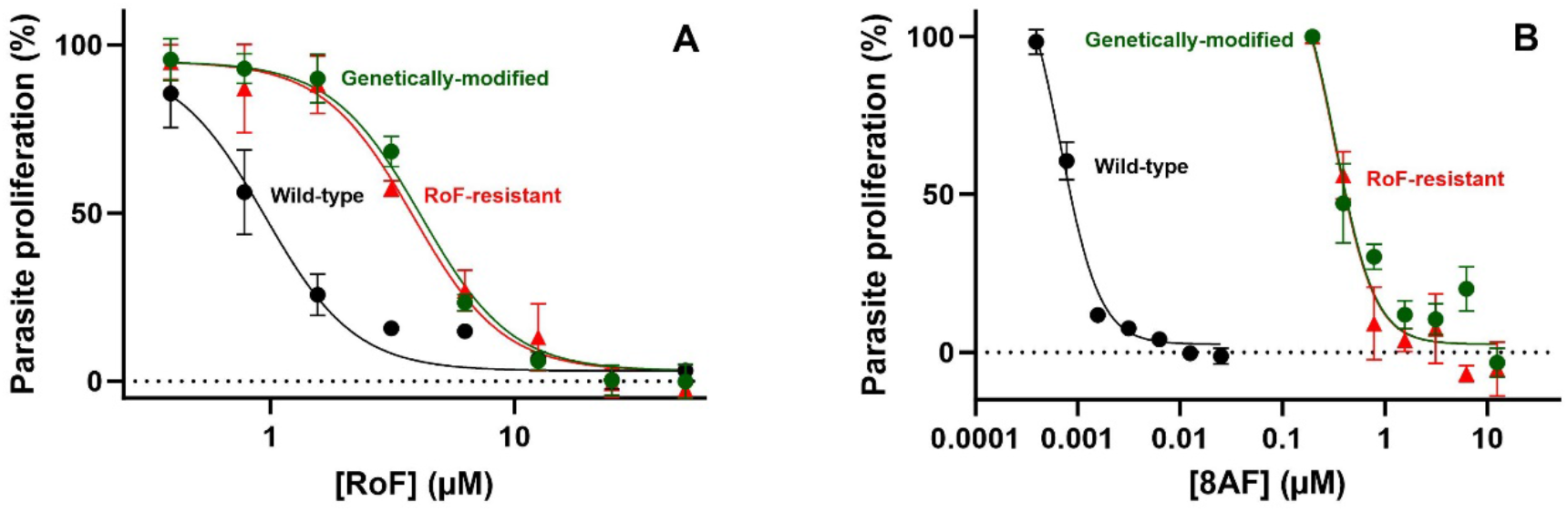
Proliferation of parent parasites (black circles), parasites selected to be RoF resistant by *in vitro* evolution (red triangles), and parasites that have had their endogenous *Pf*FK gene selectively modified (to generate the L672H mutation in the protein) by single-crossover recombination (green circles), in the presence of (A) RoF in normal RPMI-1640, (B) 8AF in riboflavin-free RPMI-1640. Values are averaged from 3-4 independent experiments, each carried out in triplicate. All error bars represent SEM and are not visible if smaller than the symbols.

### Wild-type and mutant *Pf*FK localise to the parasite cytosol

To study the localisation of the WT and mutant flavokinase we generated transgenic parasites expressing GFP-tagged versions of either the WT or mutant *Pf*FK. Lysates (from parasites that express the GFP-tagged WT and mutant forms of *Pf*FK) were analysed for protein expression using western blot. A band corresponding to a protein of the size predicted for the GFP-tagged *Pf*FK (110.7 kDa; **Figure 6A**) was observed for both parasite lines. The percentage of GFP-positive infected erythrocytes (Hoechst-positive cells) that express the WT and mutant versions of *Pf*FK was found to be approximately 25%, based on flow cytometry analysis of green GFP-fluorescence (**Figure S2**). When the parasites were examined with fluorescence microscopy, we found that both WT and mutant *Pf*FK localise to the cytosol of trophozoite and schizont stages of the parasites (**Figure 6B**).

**Figure 6:**
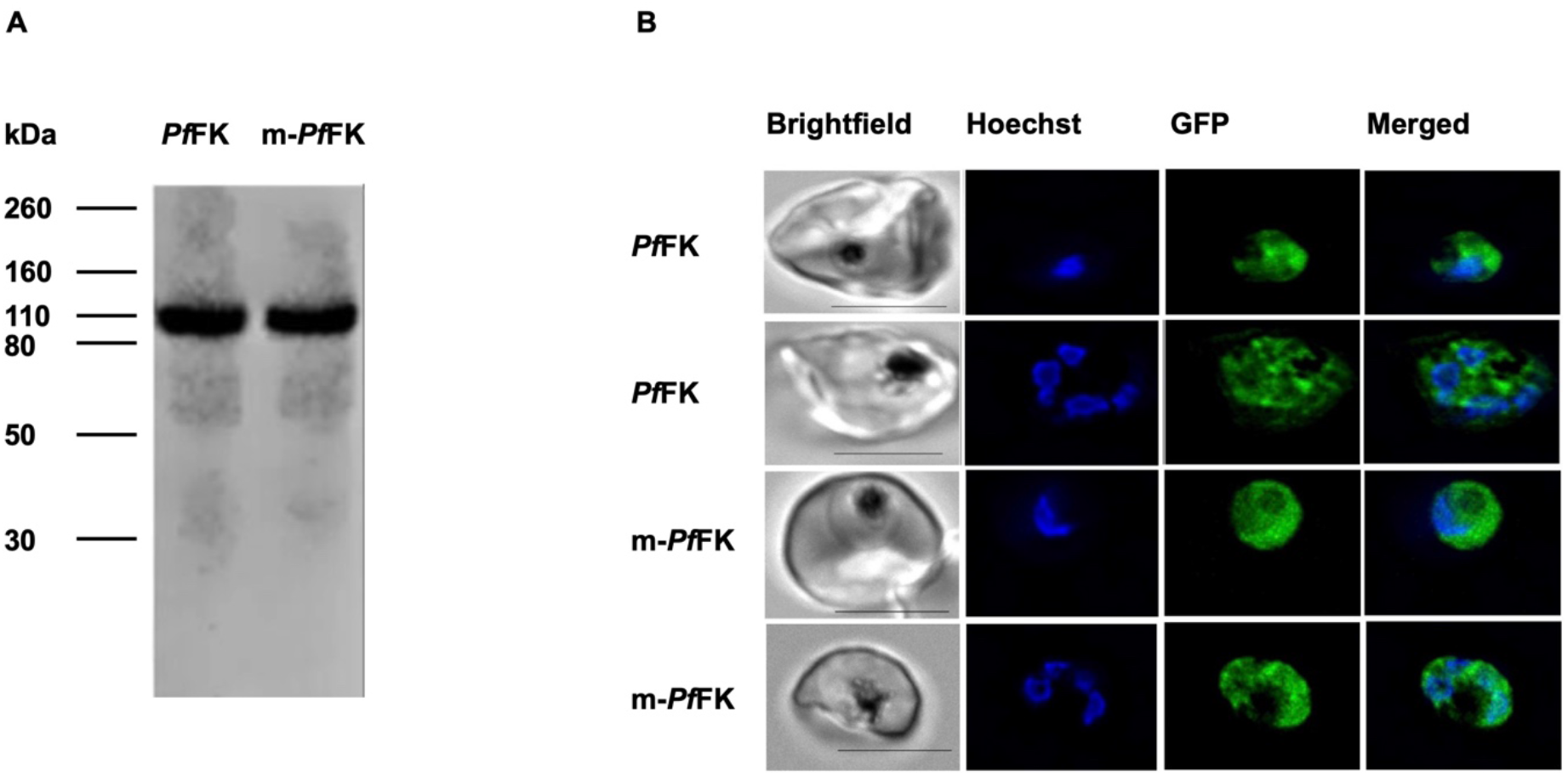
Western blot analysis (A) and confocal microscopy (B) of parasites episomally expressing wild-type and mutant *Pf*FK tagged to GFP. **(A)** Western blot analysis of protein lysates from *P. falciparum* parasites episomally-expressing wild-type *Pf*FK-GFP (left lane) or mutant *Pf*FK-GFP (m-*Pf*FK, right lane). The predicted size of *Pf*FK-GFP is 110.7 kDa. **(B)** Subcellular localization of episomally-expressed, GFP-tagged wild-type (top two rows) and mutant form (bottom two rows) of *Pf*FK (m-*Pf*FK) within trophozoite-stage and schizont-stage *P*. *falciparum-*infected erythrocytes. Brightfield images are show in the left column followed by Hoechst 33258 (DNA) labelling, GFP-fluorescence, and merged Hoechst and GFP images. Scale bars represent 4 µm.

### *Pf*FK is functional and riboflavin, RoF, and 8AF are substrates

To investigate whether *Pf*FK is responsible for the synthesis of FMN within the parasite, we purified the wild-type and mutant *Pf*FK-GFP proteins *via* immunoprecipitation using GFP-Trap^®^, carried out enzyme reactions (with the enzyme still attached to the GFP-Trap^®^, **Figure S3**) and used HPLC to detect reaction products. We found that riboflavin, RoF and 8AF are substrates of the wild-type *Pf*FK-GFP, generating FMN, roseoflavin mononucleotide (RoFMN) and 8-demethyl-8-amino-riboflavin mononucleotide (8AFMN), respectively (**Figure 7**). There was no significant difference in the initial rate of conversion of riboflavin into FMN (0.097 ± 0.012 µmol/10^10^ cells/min) and RoF into RoFMN (0.068 ± 0.009 µmol/10^10^ cells/min, P = 0.1221, unpaired, two tailed t-test, N =3) when each were individually present at 7.5 µM. On the other hand, 8AF (also present at 7.5 µM) was converted into 8AFMN at a significantly higher rate (62.5 ± 6.1 µmol/10^10^ cells/min) than both riboflavin and RoF (P = 0.0005 and 0.0005, respectively, unpaired, two tailed t-test, N = 3; **Figure 7**). We did not detect production of FAD, consistent with *Pf*FK being a monofunctional enzyme, catalysing only the conversion of riboflavin into FMN and not also acting as an FAD synthetase to produce FAD as occurs in some other organisms ^37, 38^.

**Figure 7:**
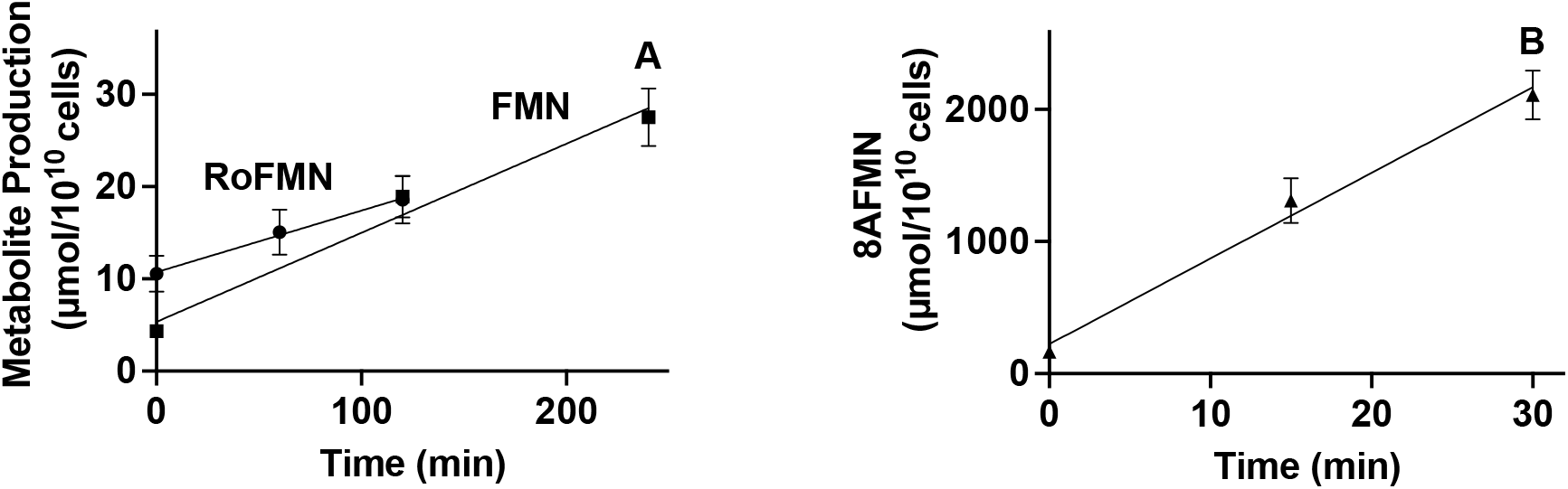
Riboflavin (A, squares), RoF (A, circles), and 8AF (B, triangles), all at 7. µM, are phosphorylated by the purified wild-type form of *Pf*FK-GFP. Values are averaged from three independent experiments, each carried out in triplicate. Error bars represent SEM and, if not visible, are smaller than the symbols.

### L672H mutation of *Pf*FK alters its affinity to its substrates

We carried out full Michaelis-Menten (MM) analysis for the conversion of riboflavin, RoF and 8AF by the WT and mutant forms of *Pf*FK (**Figure 8**). The amount of protein used was standardised across the samples using western blotting (**Figure S4**). The apparent K_M_ of the wild-type form of *Pf*FK for riboflavin was found to be 1.2 ± 0.1 µM, and not significantly different from that for RoF (1.3 ± 0.2 µM) and 8AF (1.1 ± 0.1 µM, P > 0.3, N = 3, unpaired t-test, three comparisons, N = 3, **Figure 8 and Table 1**). We also found that the V_max_ of WT *Pf*FK for riboflavin (0.12 ± 0.01 µmol/min/10^10^ cells) was not significantly different from that for RoF (0.15 ± 0.01 µmol/min/10^10^ cells, P = 0.1231, N =3, unpaired t-test, **Figure 8 and Table 1**). However, 8AF is metabolised by the wild-type form of flavokinase at much higher rate than riboflavin and RoF by three orders of magnitude (142 ± 2, P<0.0001, N = 3, unpaired t-test, **Figure 8 and Table 1**). Moreover, the V_max_/K_M_, a measurement of catalytic efficiency ^39, 40^, of *Pf*FK for 8AF is higher than those for riboflavin and RoF by a factor of 10^3^, while *Pf*FK has the same catalytic efficiency for riboflavin and RoF **(Table 1)**.

**Figure 8:**
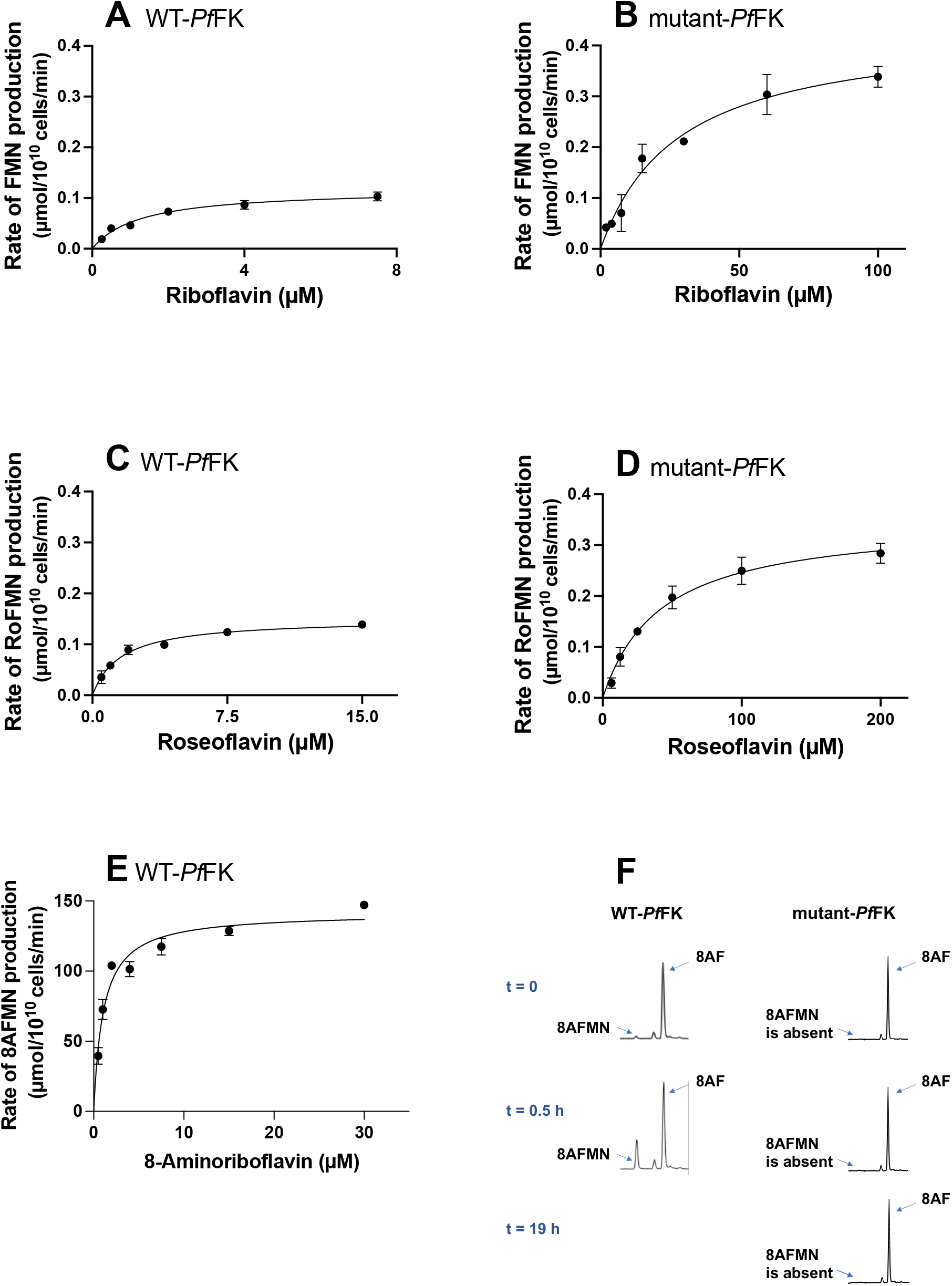
Michaelis-Menten curves for the phosphorylation of riboflavin (A and B), RoF (C and D) and 8AF (E) by the wild-type (A, C, and E) and the mutant (B, and D) flavokinase of *P.falciparum*. Panel F is an HPLC chromatogram showing the synthesis of 8AFMN from 8AF and ATP upon the addition of the wild-type *Pf*FK, while no 8AFMN production was detected upon the addition of the mutant flavokinase. Values are averaged from 3 independent experiments, each carried out in triplicate. Traces in F are representative of those obtained in 3 independent experiments and are scaled to the tallest peak. All error bars represent SEM. Error bars are not visible if smaller than the symbols.

**Table 1:** Kinetic constants of the wild-type and mutant *Pf*FK for different substrates.

| Enzyme | Substrate | $K_M$ | $V_{max}$ | $V_{max} / K_M$ |
| --- | --- | --- | --- | --- |
| | | ($\mu M$) | ($\mu mol/10^{10}$ cells/min) | (L/$10^{10}$ cells/min) |
| WT<br>*Pf*FK | Riboflavin | $1.2 \pm 0.1$ | $0.12 \pm 0.01$ | $0.10 \pm 0.01$ |
| | Roseoflavin | $1.3 \pm 0.2$ | $0.15 \pm 0.01$ | $0.12 \pm 0.01$ |
| | 8-aminoriboflavin | $1.1 \pm 0.1$ | $142 \pm 2$ | $134 \pm 7$ |
| Mutant<br>*Pf*FK | Riboflavin | $27 \pm 5$ | $0.44 \pm 0.06$ | $16 \pm 1 \times 10^{-3}$ |
| | Roseoflavin | $53 \pm 8$ | $0.35 \pm 0.03$ | $7 \pm 1 \times 10^{-3}$ |
|  | 8-aminoriboflavin | – * | – | – |

\* – not a substrate.

When MM curves were generated for *Pf*FK harbouring the L672H mutation we found that the K_M_ for riboflavin increased by a factor of 22, while the K_M_ for RoF increased by a factor of 41 **(Figure 8 and Table 1)**. This is consistent with the enzyme’s affinity for RoF being reduced by a factor of approximately 2-fold compared to riboflavin. The V_max_ values of the mutant *Pf*FK were also found to be significantly increased for riboflavin and RoF, by a factor of 3.7 and 2.3, (P = 0.002 and 0.0019, unpaired t-test, N = 3), respectively, **(Figure 8 and Table 1)**. The V_max_/K_M_ of this mutant enzyme is significantly reduced for riboflavin and RoF by a factor of 6 and 14, respectively (P = 0.007 and 0.0004 unpaired t-test, N = 3, **Table 1**). This means that the mutant *Pf*FK has a higher catalytic efficiency (by a factor of 2.3) for riboflavin than RoF. In contrast to the observations with riboflavin and RoF (with mutant *Pf*FK still accepting riboflavin and RoF, albeit with reduced affinity), 8AF is no longer a substrate of the mutant *Pf*FK, with no 8AFMN production detected even after a 19 h incubation at 37 °C (**Figure 8F**).

## Discussion

Although we have recently shown that the parasite is capable of FMN and FAD synthesis ^16^, the putative *P. falciparum* flavokinase gene remained to be experimentally verified and characterised. In this study, present data consistent with the putative *Pf*FK gene coding for a monofunctional flavokinase that utilises riboflavin, RoF and 8AF as substrates. Previous studies have shown that the flavokinase of humans and plants localises to the cell cytosol ^41, 42^. We show here that, similarly, *Pf*FK localises to the parasite cytosol. The affinity of *Pf*FK for riboflavin (K_M_ = 1.2 µM) is somewhat intermediate when compared to the reported affinity of FK for riboflavin in other organisms (K_M_ = 0.0103-180 µM) ^37, 43–46^. *Pf*FK exhibits a lower affinity for riboflavin compared to *Arabidopsis thaliana*, *Neurospora crassa*, and *Nicotiana tabacum* (Giancaspero et al., 2009, Sandoval and Roje, 2005, Rajeswari et al., 1999), while it has a higher affinity, to different degrees, when compared to the FK from organisms such as *Corynebacterium ammoniagenes*, *Megasphaera elsdenii*, *Rattus norvegicus*, *Bos taurus*, *Streptomyces davaonensis*, and *Bacillus subtilis* ^37, 38, 44, 45, 47–49^. The K_M_ of the human flavokinase for riboflavin has recently been reported to be 2.5 µM ^45^), only modestly higher than that we report here for *Pf*FK (K_M_ = 1.2 µM). However, when measured under similar conditions to those applied in our study, the K_M_ value of the human FK for riboflavin was determined to be 36 µM ^30^, a 30-fold lower affinity when compared to *Pf*FK. The higher affinity for riboflavin by *Pf*FK, when compared to the human FK, may allow the parasite to compete effectively for the riboflavin present in human serum. Perhaps a more important difference, however, at least from a therapeutic point of view, is the fact that the K_M_ values of the human FK for RoF and 8AF (160 µM and 885 µM, respectively ^30^) were substantially higher than those observed in our study for *Pf*FK (Table 1). This represent a much bigger difference in affinity between the parasite and host FK (100-800-fold), clearly demonstrating that the active site of *Pf*FK can be selectively targeted by riboflavin analogues.

The observation that the L672H mutation in *Pf*FK confers resistance against RoF and 8AF (**Figure 2**) indicates that this mutation likely affects the ability of *Pf*FK to bind to these two riboflavin analogues. Consistent with this is the demonstration that the L672H mutation modulates *Pf*FK activity and substrate specificity (**Figure 8**), and with L672’s predicted location in the riboflavin binding site (**Figure 3**), which is likely to be shared with the two analogues.

We previously found that RoF and 8AF can kill *P. falciparum* parasites at nanomolar concentrations in medium containing riboflavin levels comparable to those found in human plasma ^16^. The fact that increasing the extracellular riboflavin concentration reduced the antiplasmodial potency of RoF and 8AF, together with our observation that parasite FMN production is compromised by RoF and 8AF, is consistent with these compounds interfering with riboflavin metabolism and/or utilisation, either as inhibitors or substrates of the enzymes involved ^16^. One possibility, therefore, is that RoF and 8AF kill *P. falciparum* parasites by competing with riboflavin, reducing in the formation of FMN and FAD, inactivating the downstream flavoenzymes that rely on these flavins. In this study, we show that RoF and 8AF are substrates for *Pf*FK, generating RoFMN and 8AFMN, respectively. Another possibility is that the metabolites generated from RoF and 8AF, RoFMN and 8AFMN, respectively, are toxic to the parasite via inhibition of FMN-utilising enzymes, such as dihydroorotate dehydrogenase^50, 51^. These two possible mechanisms are almost certainly working in tandem to inhibit parasite proliferation.

Any generation of FAD antimetabolites from RoFMN and 8AFMN by FAD synthetase would also inactivate FAD-utilising enzymes, such as glutathione reductase, thioredoxin-disulfide reductase and pyridoxine 5-dehydrogenase ^52–54^. Of course, a highly likely possibility is that both mechanisms (i.e. the reduction of FMN and FAD as well as the generation of their antimetabolites) are contributing to the antiplasmodial effect of these compounds. A decrease in the levels of FMN and FAD would boost the performance of RoFMN and 8AFMN, even though these reduced levels alone may not be adequate to eliminate the parasite on their own. In support of this possibility is our observation that although 8AF is not a substrate of mutant *Pf*FK (Fig. 8F), its antiplasmodial activity is antagonised by increasing the extracellular concentration of riboflavin (**Figure 2B**, **Figure 4B**). This is consistent with the activity of 8AF, at least against RoF-resistant parasites, being due to a reduction in FMN synthesis by competitively (with riboflavin) inhibiting m-*Pf*FK. Complete characterisation of the other potential enzymes involved in the activity of RoF and 8AF (such as FAD synthetase) of the parasites and human, will allow for the identification of regulatory and kinetic differences, which could facilitate the development of targeted inhibitors.

## Acknowledgements

We are grateful to The Australian Red Cross Lifeblood Canberra Branch for the provision of red blood cells. AH received funding from the Australian Government’s Research Training Program. This work was supported by the Alliance Berlin-Canberra “Crossing Boundaries: Molecular Interactions in Malaria”, which is co-funded by the Australian National University and a grant from the Deutsche Forschungsgemeinschaft (DFG) for the International Research Training Group (IRTG) 2290.

---

**Figure S1:**
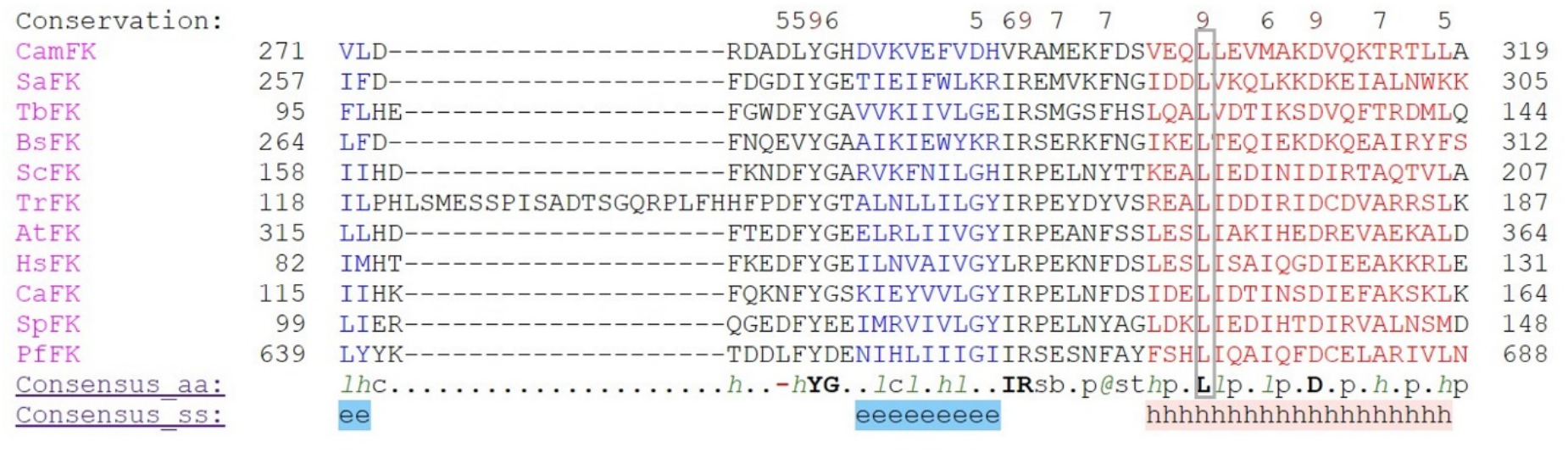
The alignment of residues 639-688 of *Pf*FK with the relevant sections of the flavokinase proteins from other organisms. The flavokinase (FK) homologues (accession numbers in brackets) from the following species were aligned: Pf: *P. falciparum* (Q8IDB3); Hs: *Homo sapiens* (Q969G6); Sp: *Schizosaccharomyces pombe* (O74866); Bs: *Bacillus subtilis* (P54575); Tb: *Trypanosoma brucei* (Q38DG4); At: *Arabidopsis thaliana* (Q84MD8); Sa: *Streptococcus agalactiae serotype* Ⅲ (Q8E5J7); Ca: *Candida albicans* (Q5A015); Tr: *Trichophyton rubrum* (F2SJS4); Sc: *Saccharomyces cerevisiae* (Q03778); Cam: *Corynebacterium ammoniagenes* (Q59263). L672 of *Pf*FK and the corresponding leucine in the flavokinase homolouges of other organims is marked with a grey rectangle and is conserved in all of the 10 sequences. Values at and above the conservation index cut-off (5) are displayed above the amino acid. Consensus aa refers to the consensus level alignment parameters for the consensus amino acid sequence. This is displayed if the weighted frequency of a certain class of residues in a position is above 0.8. Consensus symbols are as follows: conserved amino acids: bold and uppercase letters; aliphatic (I, V, L): l; aromatic (Y, H, W, F): @; hydrophobic (W, F, Y, M, L, I, V, A, C, T, H): h; alcohol (S, T): o; polar residues (D, E, H, K, N, Q, R, S, T): p; tiny (A, G, C, S): t; small (A, G, C, S, V, N, D, T, P): s; bulky residues (E, F, I, K, L, M, Q, R, W, Y): b; positively charged (K, R, H): +; negatively charged (D, E): -; charged (D, E, K, R, H): c. Elements of secondary structure (ss) are indicated: h = alpha helix, e = beta strand.

**Figure S2:**
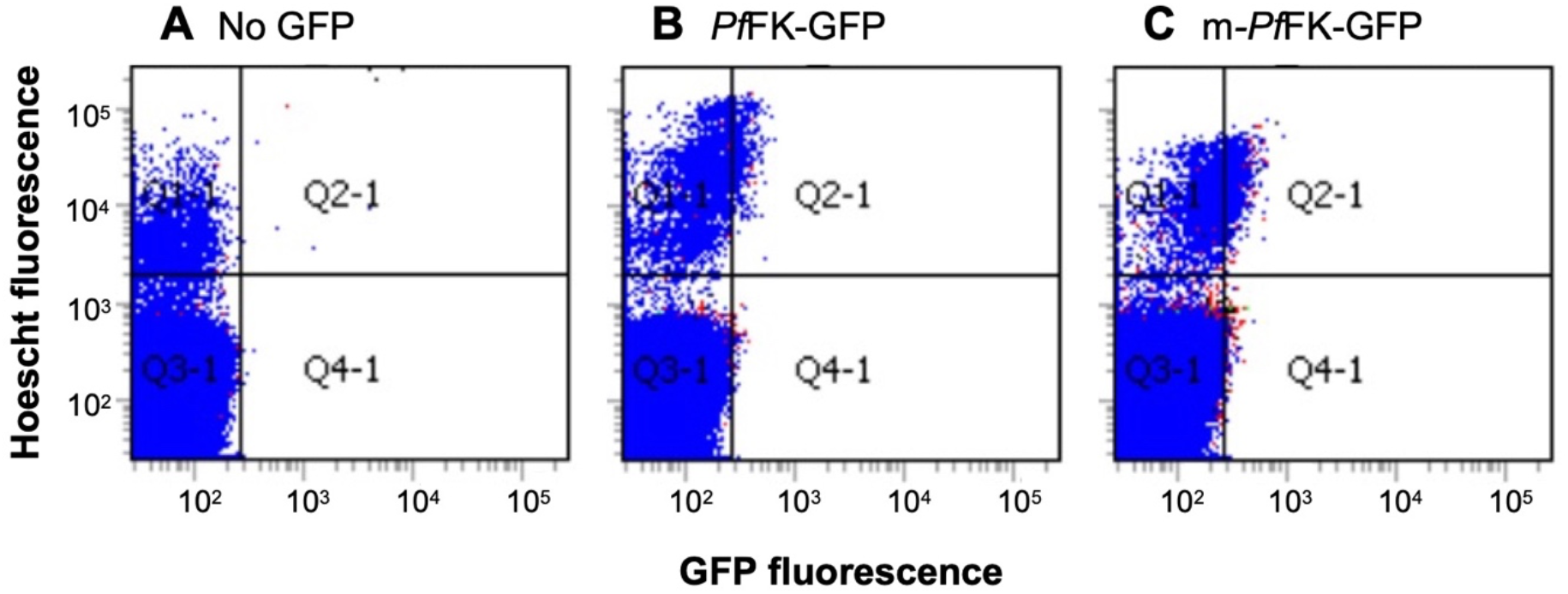
Determination of the proportion of trophozoite-stage parasites expressing the WT-*Pf*FK-GFP and mutant-*Pf*FK-GFP. The forward scatter intensity on each x-axis corresponds to GFP fluorescence and the y-axis corresponds to the intensity of Hoescht fluorescence (to identify infected erythroctyes). Using 3D7 trophozoites to create a gating threshold below which parasites were considered as auto-fluorescent, the proportion of GFP-positive cells were estimated in each transgenic line (B: WT-*Pf*FK-GFP (27 ± 6% n = 3) and C: mutant-*P*fFK-GFP (27 ± 7% n = 3)). Data is representative of those obtained from three independent experiments.

**Figure S3:**
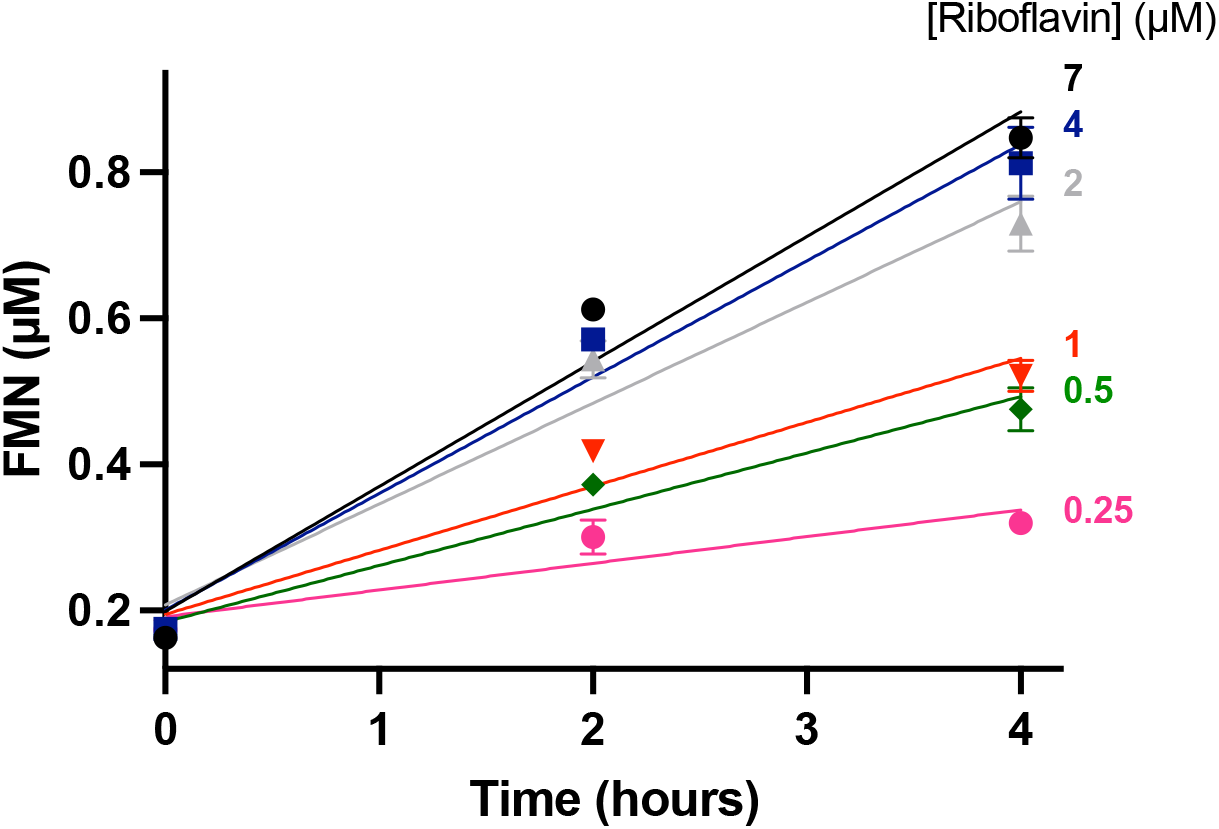
Formation of FMN from riboflavin and ATP by purified *Pf*FK-GFP. The rate of FMN production increased as the concentration of riboflavin was increased from 0.25 to 7 µM. Values are averaged from three independent experiments, each carried out in triplicate. Error bars represent SEM and are not visible if smaller than the symbols.

**Figure S4:**
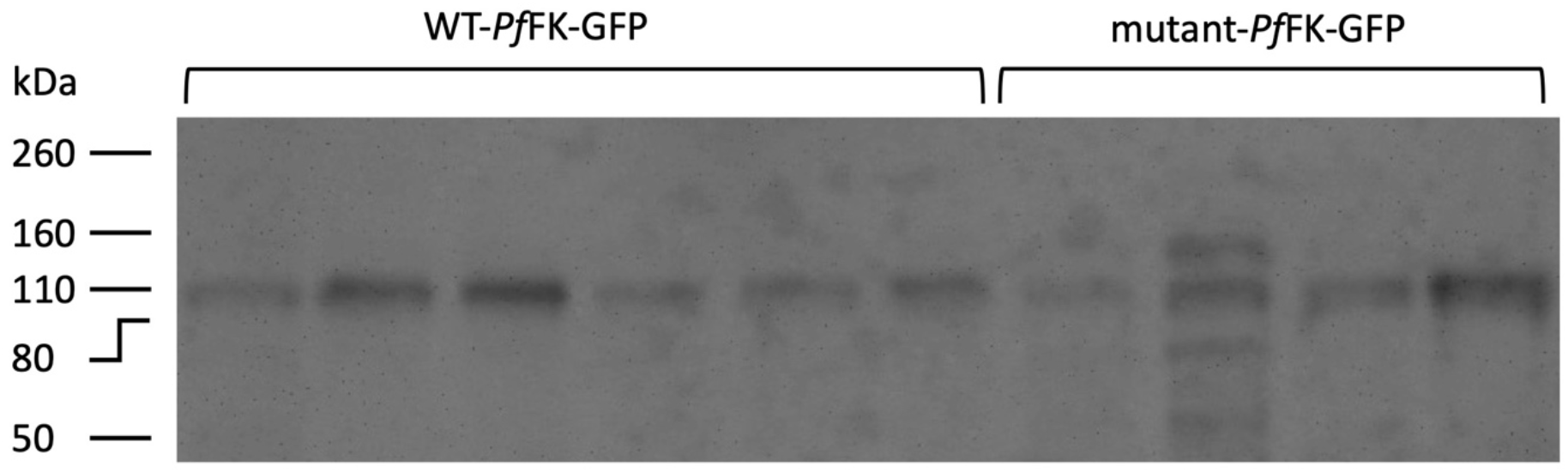
Western blot analysis of WT-*Pf*FK-GFP, and mutant-*Pf*FK-GFP in GFP-Trap-immunoprecipitated preparations. Purified *Pf*FK samples, each prepared from 1 × 10^10^ isolated *P. falciparum* trophozoites expressing WT-*Pf*FK-GFP (Lane 1 to 6; riboflavin was tested using the first three samples (lanes 1-3) while RoF and 8AF were tested, in the same experiment using the next three samples (lanes 4-6)) or mutant-*Pf*FK-GFP (Lane 7-10; each sample used to test riboflavin, RoF and 8AF) were analysed by western blotting. Image J software was used to quantify the band intensity and were then standardized to the lowest value in preparation for the enzyme activty assays.

**Table S1:** Key sequencing results from whole genome sequencing of WT and RoF-resistant parasites.

| Gene Name | Gene Description | Effect | Codon Change | Amino Acid Change |
| --- | --- | --- | --- | --- |
| **RoF-A-Clone-E8** |  |  |  |  |
| PF3D7_0614200 | Conserved *Plasmodium* protein unknown function | Non-synonymous | Gat/Aat | D859N |
| PF3D7_1359100 | Riboflavin kinase FAD synthase family protein putative | Non-synonymous | cTt/cAt | L672H |
| PF3D7_1038400 | Gametocyte-specific protein | Non-synonymous | Ata/Gta | I8312V |
| PF3D7_1038400 | Gametocyte-specific protein | Non-synonymous | gaA/gaC | E8342D |
| PFC10_API0033 | Apicoplast ribosomal protein S5 | Non-synonymous | aGa/aAa | R118K |
| **RoF-A-Clone-B6** |  |  |  |  |
| PF3D7_1410300 | Conserved *Plasmodium* protein unknown function | Non-synonymous | Gaa/Aaa | E4170K |
| PF3D7_1471900 | Conserved *Plasmodium* protein unknown function | Non-synonymous | aaT/aaA | N907K |
| PF3D7_1303800 | Conserved *Plasmodium* protein unknown function | Non-synonymous | Caa/Aaa | Q418K |
| PF3D7_1359100 | Riboflavin kinase FAD synthase family protein putative | Non-synonymous | cTt/cAt | L672H |
| PFC10_API0033 | Apicoplast ribosomal protein S5 | Non-synonymous | aGa/aAa | R118K |
| **RoF-B-Clone-E11** |  |  |  |  |
| PF3D7_1420000 | Splicing factor subunit putative | Non-synonymous | tCa/tTa | S227L |
| PF3D7_0728100 | Conserved *Plasmodium* membrane protein unknown function | Non-synonymous | aaT/aaA | N5064K |
| PF3D7_1359100 | Riboflavin kinase FAD synthase family protein putative | Non-synonymous | cTt/cAt | L672H |
| PFC10_API0022 | Probably protein unknown function | Non-synonymous | tCa/tTa | S4L |
| PFC10_API0033 | Apicoplast ribosomal protein S5 | Non-synonymous | aGa/aAa | R118K |
| **RoF-B-Clone-G2** |  |  |  |  |
| PF3D7_0406500 | Conserved *Plasmodium* protein unknown function | Non-synonymous | Ctt/Att | L1137I |
| PF3D7_1316400 | Conserved *Plasmodium* protein unknown function | Non-synonymous | aaC/aaA | N2K |
| PF3D7_1359100 | Riboflavin kinase FAD synthase family protein putative | Non-synonymous | cTt/cAt | L672H |
| PFC10_API0022 | Probably protein unknown function | Non-synonymous | Caa/Aaa | Q15K |
| PFC10_API0022 | Probably protein unknown function | Non-synonymous | tCa/tTa | S4L |
| PFC10_API0033 | Apicoplast ribosomal protein S5 | Non-synonymous | aGa/aAa | R118K |
| PFC10_API0055 | Apicoplast ribosomal protein S4 | Non-synonymous | tGt/tAt | C31Y |
| PFC10_API0055 | Apicoplast ribosomal protein S4 | Non-synonymous | Ggt/Agt | G25S |

